# Cholesterol links blood feeding to mosquito development and reproduction

**DOI:** 10.64898/2026.05.15.725405

**Authors:** Sagiv Cohen, Benjamin Trabelcy, Doron Shalom Yishai Zaada, Philippos Aris Papathanos, Yoram Gerchman, Alon Silberbush, Amir Sapir

## Abstract

Mosquitoes undergo major metabolic remodeling as they transition from aquatic larvae to terrestrial, blood-feeding adults, yet the biochemical principles supporting these life cycle transitions remain poorly understood^1^. Early metabolic studies established that mosquitoes are sterol auxotrophs^2,3^; however, the extent to which this auxotrophy influences developmental progression and stage transitions remains unknown. Here, using sterol-defined culture systems, we uncover a central and stage-specific role for sterol metabolism in mosquitoes. We show that eggs of multiple mosquito species, from *Anopheles gambiae* (Giles 1902; hereafter *A. gambiae*) to *Aedes* species in both wild and laboratory settings, are enriched in cholesterol consistent with maternal provisioning. In the Asian tiger mosquito *Aedes albopictus* (Skuse, 1894; hereafter *Ae. albopictus*), egg cholesterol supports early larval development, after which larvae depend on the acquisition of dietary sterols to complete development. Like cholesterol, dietary plant- and fungal-derived sterols support late larval development; however, their utilization is associated with the accumulation of the intermediate sterol desmosterol. Using biochemical assays, we showed that desmosterol is converted to cholesterol through the activity of the *Ae. albopictus* DHCR-24 enzyme. This metabolic axis of desmosterol-to-cholesterol conversion supports late larval development when mosquitoes rely on plant- and fungal-derived sterols for development. *dhcr-24* expression is upregulated during the developmental window of dietary sterol acquisition and is selectively induced by its substrate desmosterol, revealing a diet-encoded regulatory mechanism that coordinates metabolic conversion capacity. Extending the analysis to adulthood, we developed a sterol-defined artificial blood-feeding system that enables precise manipulation of dietary sterols in reproductive females and show that cholesterol availability in the diet of females is an essential metabolic driver of egg laying. These findings reveal a developmentally coordinated sterol-use strategy in the life cycle of *Aedes* mosquitoes, identifying sterol utilization as a central physiological axis linking development, blood feeding, and reproduction, and revealing a sterol-dependent metabolic vulnerability in mosquitoes.

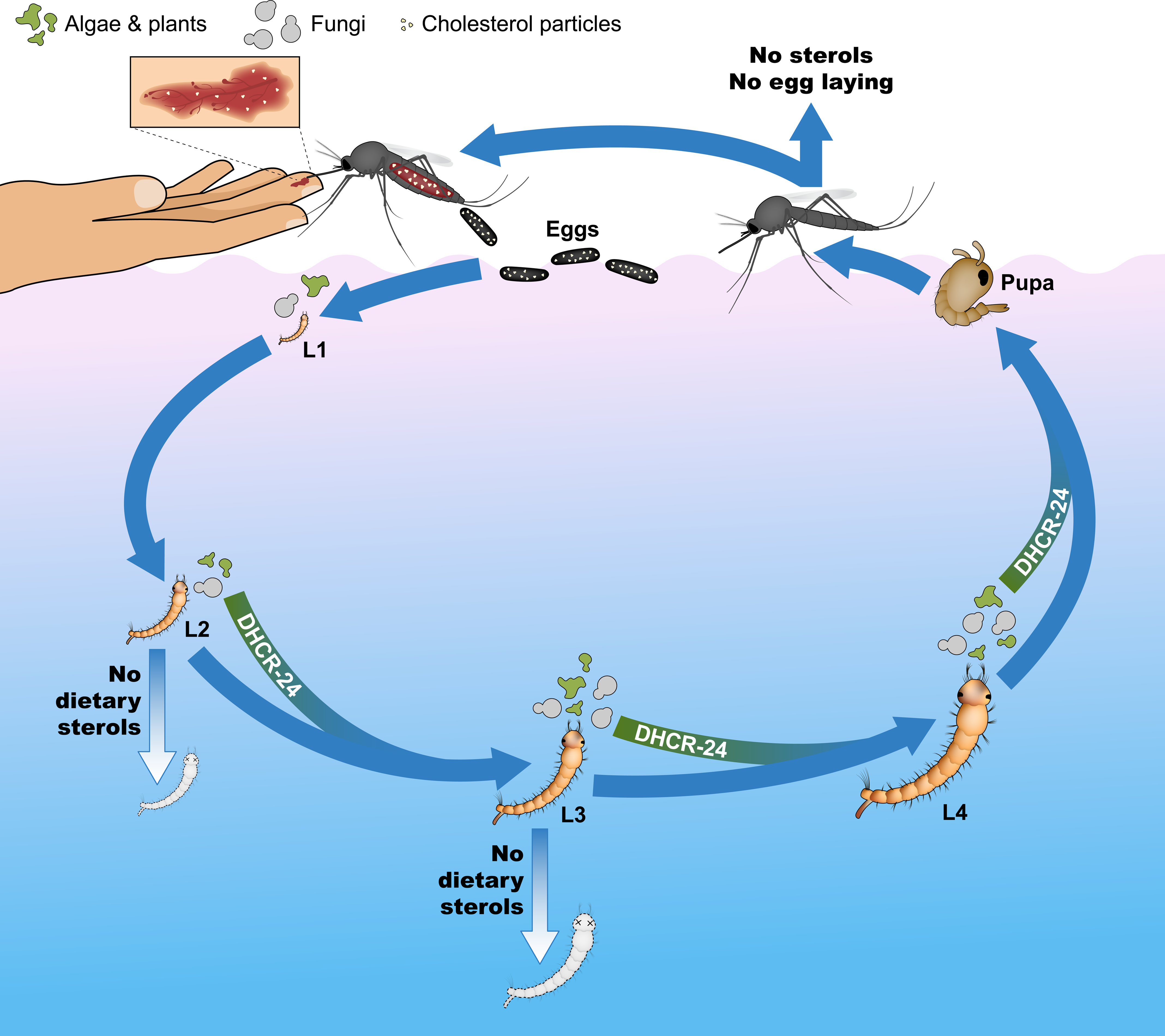

## Introduction

Sterols are essential components of eukaryotes, serving as structural elements of membranes and as precursors for diverse signaling molecules^4,5^. Unlike vertebrates, most invertebrates, including insects, lack the capacity for *de novo* sterol synthesis, relying instead on the dietary acquisition of sterols as essential nutrients to sustain development, reproduction, and viability^6–9^. Despite this sterol auxotrophy, how different insects acquire, process, and deploy sterols across their life cycles remains only partially understood^10^. Mosquitoes provide a well-studied model to investigate how nutrients are acquired and utilized in insects, as dietary requirements differ across life stages and sexes. For example, adult males and females consume nectar, whereas females additionally require blood feeding to support reproduction^11^. Early work has shown that two species of mosquitoes, *Aedes aegypti* (Linnaeus, 1762; hereafter *Ae. aegypti*)^2,12,13^ and *Culex pipiens* (Linnaeus, 1758; hereafter *C. pipiens*)^14^, depend on dietary cholesterol for completion of their life cycle. Subsequent studies demonstrated that multiple plant sterols, including β-sitosterol, stigmasterol, and campesterol, as well as the major fungal sterol ergosterol, can support the development of *C. pipiens* larvae to the adult stage^3^. Metabolic labeling experiments further support this observation, demonstrating the capacity of *Ae. aegypti* to convert plant-derived sterols into cholesterol^15^, which in most insects is the precursor for the molting hormone ecdysone^16,17^. Despite the importance of this axis for the mosquito life cycle, the genetic and metabolic mechanisms governing sterol acquisition, conversion, and utilization across development remain incompletely understood.

Here, we first profile the composition and levels of sterols in eggs from multiple mosquito species and then use *Ae. albopictus* as a model to establish a life cycle–resolved analysis of sterol metabolism in mosquitoes. By integrating sterol-defined culture systems, biochemical analyses, gene expression profiling, and reproductive assays, we uncover a developmentally and metabolically regulated sterol utilization program. This program mechanistically links environmental sterol utilization during larval development with cholesterol acquisition through blood feeding in adult females, which is essential for efficient egg laying and reproduction.

## Results

To characterize the identity, abundance, and developmental dynamics of sterols in mosquitoes, we first asked whether and which sterols are present in mosquito eggs, as their sterol composition has not been previously characterized. To capture a broad range of ecological and reproductive strategies, we analyzed eggs of mosquitoes from the wild that lay eggs in rafts, and from laboratory-reared mosquitoes maintained on controlled blood diets that lay individual eggs. Using gas chromatography–mass spectrometry (GC–MS), we found that cholesterol (Fig. S1A) is abundant in eggs across species, spanning a broad range of cholesterol content per egg, from ∼4.5 ng in *A. gambiae* to ∼83 ng in *Ae. aegypti* (Fig. 1A; Table S1). For example, *Ae. albopictus* eggs contain an average of 31.8 ng cholesterol per egg, corresponding to approximately 0.4% of egg mass, based on reported egg mass values^18^. Apart from cholesterol (Fig. 1B), no other sterols were detected in any egg samples above the GC–MS detection limit, supporting free cholesterol as the predominant egg sterol. To investigate whether egg cholesterol contributes to embryonic and early larval development, we focused subsequent experiments on *Ae. albopictus*, reflecting the public-health relevance of *Aedes* arbovirus vectors while providing a tractable foundation for cross-species comparisons. To determine the fate and function of egg cholesterol, we established a sterol-defined larval culture system maintained under sterol-free conditions, in which larvae were individually reared with a diet containing exclusively *E. coli* as a food source, which, like most bacteria, neither synthesizes nor modifies sterols^19^. GC-MS analysis revealed that, under sterol-free conditions, cholesterol progressively declined during *Ae. albopictus* larval development (Figs. 1C, D, S1B). To assess the biological significance of this decline, we examined larval survival under sterol-free conditions and found that larvae displayed progressive mortality beginning at the L3 stage (Fig. 1E), coinciding with depletion of egg cholesterol. We therefore tested whether dietary cholesterol could rescue larval lethality and support larval development. Based on previous work in *C. pipiens* demonstrating robust support of larval development at 13 µM cholesterol^3^, we used the same concentration in our experiments. Dietary cholesterol supplementation rescued larval lethality, allowing individuals to complete larval development, pupate, and emerge as adults at high rates (Fig. 1E). These findings demonstrate that dietary sterol acquisition during larval development is required for developmental progression from late larval stages to adulthood. Consistent with single-larva experiments, dietary cholesterol was also required for survival when larvae were reared in populations (Fig. 1F), a condition that more closely reflects the natural ecology of mosquito larvae, which develop in dense cohorts where nutrient availability constrains developmental success. To assess the developmental and morphological effects of sterol deprivation, we examined individually cultured larvae across developmental time points under cholesterol-depleted and supplemented conditions. Sterol-deprived larvae were smaller and paler than cholesterol-supplemented controls (Fig. 1G), without overt morphological abnormalities. Next, we examined the morphology of live and dead larvae from the sterol-free culture. Whereas live sterol-deprived larvae were significantly smaller than cholesterol-replete larvae at days 8 and 10 post-feed (Fig. 1H), larvae that died in sterol-free culture showed reduced size and pigmentation (Fig. S1C). Biochemical profiling by GC–MS revealed that dietary cholesterol supplementation markedly increased the levels of larval cholesterol and its immediate byproduct 7-Dehydrocholesterol (7-DHC) beginning at the L3 stage (Figs. 1I, J; S1D, E). Notably, earlier stages contained cholesterol levels comparable to those detected in eggs (Fig. 1D), consistent with an essential role for dietary sterol acquisition in supporting late larval development.

**Figure 1.**
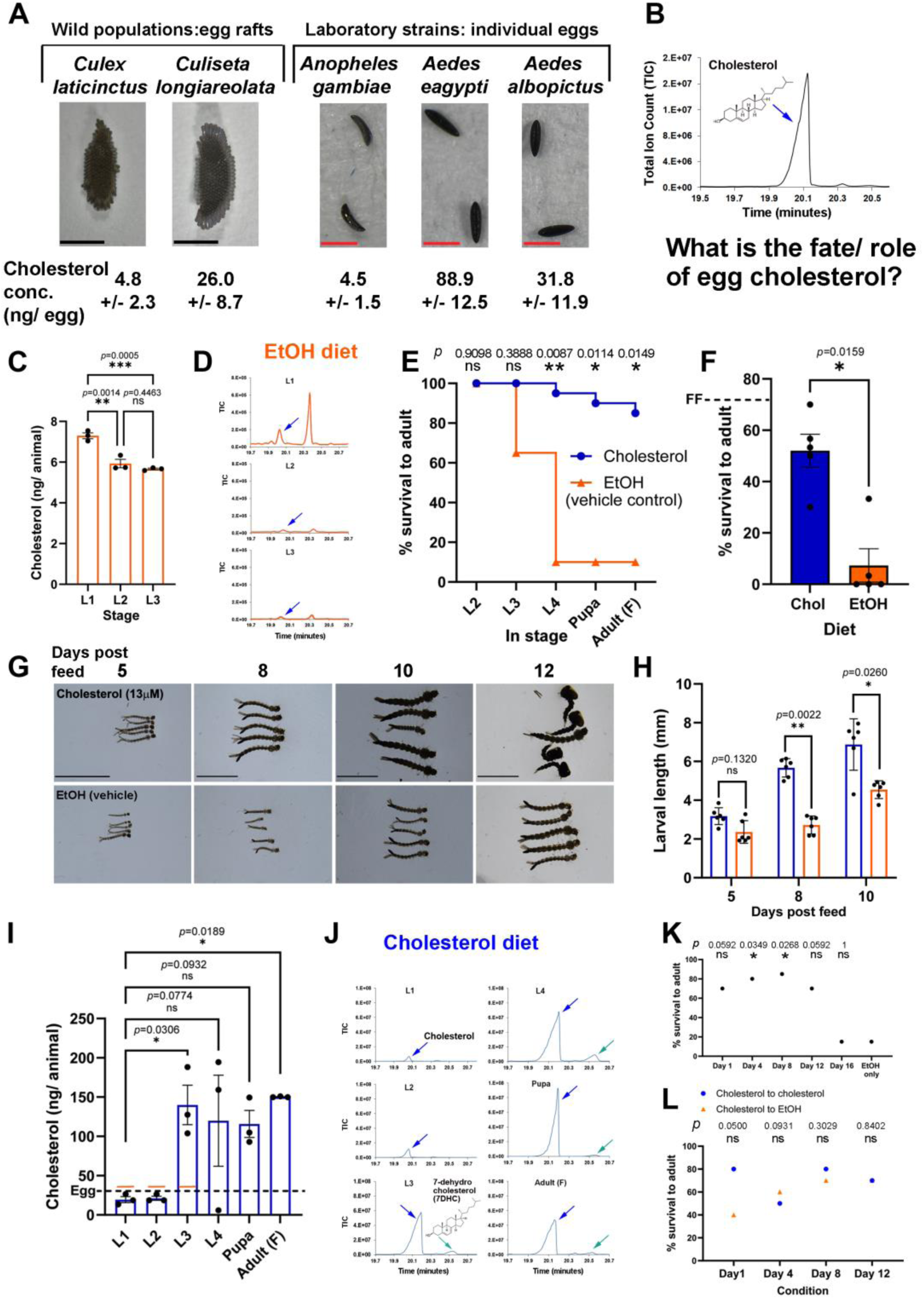
Cholesterol availability is essential for *Ae. albopictus* larval development and survival. **A.** Images of eggs from different mosquito species and their cholesterol content quantified using a calibration curve (see Methods). Scale bars, 1 mm in images of rafts and 250 µm in images of individual eggs. **B.** Representative GC–MS chromatogram of sterols from *Ae. albopictus* eggs identified using reference libraries and diagnostic ions. **C.** Cholesterol content per animal in vehicle ethanol (EtOH)-only cultures. n = 3 biological replicates of 20 larvae; one-way ANOVA test limited to L1 until L3 due to lack of survival at later stages, followed by Tukey’s post hoc test. Henceforth, normality was assessed using the Shapiro–Wilk test and equality of variance using the F test; error bars represent SEM. **D.** Representative GC–MS chromatograms from EtOH-only cultures. Hereafter, the sterol region of the chromatograms (19.7-21 minutes) is presented. **E.** Survival from L2 to adulthood under cholesterol-supplemented or ethanol (EtOH; vehicle) conditions; n = 20 individually reared larvae; Chi-square test. Hereafter, at the adult stage, only females, labeled as Adult (F), were analyzed. **F.** Survival to adulthood in population-based experiments; n = 30 larvae across 10 cups; unpaired two-tailed Mann–Whitney test. The dashed “FF” line indicates the average percent survival across all developmental stages for larval populations reared on a standard fish food diet. **G.** Representative images of larvae grown individually in single-larva cultures; anterior to the left; scale bars, 5 mm. **H.** Larval length (mm); n = 6 larvae; unpaired two-tailed Mann–Whitney test. **I.** Cholesterol content per animal across developmental stages in cholesterol-supplemented cultures. n = 3 biological replicates of pooled 20 larvae; one-way ANOVA test followed by Dunnett’s post hoc test. **J.** Representative GC–MS chromatograms from 20 cholesterol-supplemented larvae across developmental stages. **K.** Survival of larvae grown in sterol-free culture and supplemented with cholesterol at defined time points; n = 20 individually reared larvae; Chi-square test. **L.** Cholesterol-to-EtOH transfer experiment, with cholesterol-to-cholesterol transition as control. n = 3 independent cultures per condition, 10 larvae per culture; Chi-square test.

Having established that cholesterol supplementation increases larval sterol levels and rescues the developmental arrest under sterol-free conditions, we next sought to define the timing and dynamics of this dietary requirement. To this end, larvae were cultured under sterol-free conditions and supplemented with cholesterol at progressively later developmental stages. Each time point of cholesterol supplementation was compared to a single sterol-free control group that was maintained without sterols throughout the experiment. We identified a permissive window of approximately eight days during which larvae survive in the absence of sterols without compromising survival to adulthood upon cholesterol reintroduction (Fig. 1K). To define the temporal window during which dietary cholesterol can rescue the effects of sterol depletion, larvae were initially maintained under sterol-free conditions and then transferred to cholesterol-supplemented or EtOH control cultures at 0, 4, 8, or 12 days after hatching. Exposure limited to the first day of development was insufficient to rescue survival to adulthood, as reflected by the marked difference in the fraction of adults between cholesterol- and EtOH-treated groups (Fig. 1L). In contrast, prolonged cholesterol exposure resulted in similar adult fractions in the two conditions following transfer. Notably, rescue was already evident after four days of cholesterol availability, corresponding to the L2 stage, suggesting that dietary cholesterol can support larval development at early stages even though the associated increase in larval cholesterol levels was not detectable by GC–MS.

Collectively, these data confirm that *Ae. albopictus* is a sterol auxotroph and reveal a developmental transition from reliance on egg cholesterol in early larval stages to the acquisition of dietary sterols at later stages to complete its life cycle.

Mosquito larval habitats are suggested to contain diverse environmental sterols^20^, including the plant sterols β-sitosterol, stigmasterol, and campesterol, as well as the major fungal sterol ergosterol. Building on previous studies showing that dietary plant and fungal sterols can support larval development in *Ae. aegypti*^2^ and *C. pipiens*^3^, we first tested whether these environmental sterols can similarly support *Ae. albopictus* development. Using the sterol-defined system we developed, we tested whether *Ae. albopictus* larvae can develop to adulthood when supplemented with the plant sterol stigmasterol and the fungal sterol ergosterol, two sterols synthesized by members of multiple major algal phyla^21^. Algae constitute one of the major natural food sources for mosquito larvae^22^, making these sterols environmentally relevant candidates to support development. In contrast to sterol-free conditions, supplementation with either stigmasterol or ergosterol significantly supported progression beyond early instars and enabled completion of development to adulthood in mosquitoes reared as single animals (Figs. 2A; S2A) and as populations (Fig. 2B). These findings demonstrate that *Ae. albopictus* larvae can utilize plant- and fungal-derived sterols to complete development. Consistent with their ability to support survival to adulthood, larvae supplemented with stigmasterol or ergosterol displayed morphology comparable to cholesterol-supplemented controls (Fig. 2C). Average body length was also similar across most developmental stages (Fig. 2D), except for ergosterol-fed larvae at day 8. Because stigmasterol supports survival and growth more robustly than ergosterol (Fig. 2A, C), it was used in the following experiments.

**Figure 2.**
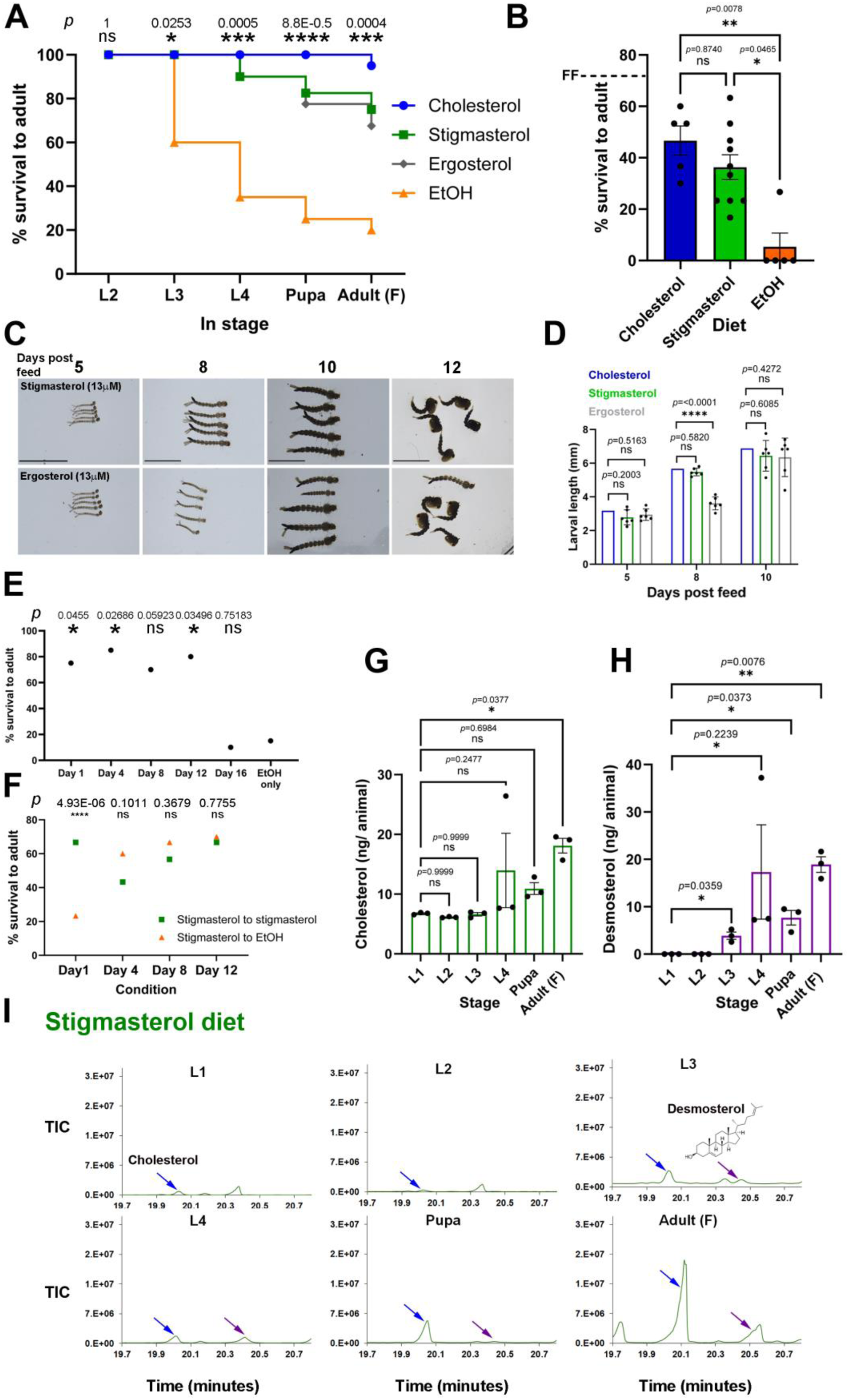
Stigmasterol supplementation supports Ae. albopictus development and results in desmosterol accumulation. **A.** Survival at each developmental stage (L2–adult) under cholesterol, stigmasterol, ergosterol, or EtOH conditions; n = 40 individually reared larvae per condition for stigmasterol and ergosterol and n = 20 individually reared larvae per condition for cholesterol and EtOH; Chi-square test results for stigmasterol in comparison to EtOH control. Statistical analysis of all sterols is shown in Fig. S2A. **B.** Survival to adulthood in a population-based experiment under supplemented sterol diets; n = 30 larvae across 10 cups for stigmasterol and n = 30 larvae across 5 cups for cholesterol and EtOH controls; Kruskal-Wallis test followed by Dunn’s post hoc test. The dashed “FF” line indicates the average percent survival across all developmental stages for larval populations reared on a standard fish food diet. **C.** Representative images of larvae grown individually in single-larva cultures under stigmasterol or ergosterol supplementation; anterior to the left; scale bars, 5 mm. **D.** Larval length (mm); n = 6 larvae; one-way ANOVA test followed by Dunnett’s post hoc test for stigmasterol and cholesterol comparison, and Kruskal–Wallis test followed by Dunn’s post hoc test for ergosterol and cholesterol comparison. **E.** Survival to adulthood following stigmasterol supplementation initiated at different developmental days; n = 20 individually reared larvae per condition; Chi-square test. **F.** Stigmasterol-to-ethanol transition experiment with cholesterol-to-cholesterol control; n = 3 cultures per condition, 10 larvae per culture; Chi-square test. **G.** Cholesterol content per animal across developmental stages in stigmasterol-supplemented cultures; n = 3 biological replicates of pooled 20 larvae; one-way ANOVA test followed by Dunnett’s post hoc test. **H.** Desmosterol content per animal across developmental stages in stigmasterol-supplemented cultures; n = 3 biological replicates of pooled 20 larvae; one-sample Student’s t-test against zero, as desmosterol was undetectable at L1. **I.** Representative GC–MS sterol chromatograms from 20 larvae reared on a stigmasterol-supplemented diet across developmental stages.

To assess the temporal window of stigmasterol utilization, we timed its supplementation following growth in sterol-free conditions. *Ae. albopictus* larvae retained the capacity to recover and complete development after a 12-day window of sterol deprivation (Fig. 2E), demonstrating resilience to sterol-limited conditions likely encountered in larval habitats. Whereas removal from stigmasterol on day one impaired development, removal of stigmasterol at later time points had little effect on adult survival (Fig. 2F), indicating that accumulated stigmasterol is sufficient to support subsequent development. The capacity of environmental sterols to support *Ae. albopictus* development raises the possibility that these nutrients are converted into cholesterol and downstream sterols. Using calibration curves to quantify sterol levels (Fig. S2B), GC–MS analysis revealed that larvae reared on stigmasterol as the sole dietary sterol accumulated increasing levels of cholesterol as development progressed, with a marked rise from the L3 stage onward (Figs. 2G, S2C, D). To test whether cholesterol detected in larvae could arise from the conversion of plant and fungal sterols by the bacterial feed, we performed GC–MS analysis of sterol-defined cultures containing *E. coli* without mosquitoes. At the end of the experiment, lipid profiles matched the supplied sterols, with no detectable bacterial conversion of plant- or fungal-derived sterols to cholesterol (Fig. S2E). Although contributions from the larval microbiota cannot be excluded, these results rule out sterol synthesis or modification by the *E. coli* feed, indicating that cholesterol detected in larvae beyond stage two arises from mosquito-intrinsic conversion of stigmasterol to cholesterol.

Together, these results show that *Ae. albopictus* larvae can exploit plant- and fungal-derived sterols, likely among the predominant sterol sources in natural aquatic habitats, to support growth and development.

GC–MS analysis of larvae fed stigmasterol (Fig. S2C) or ergosterol (Fig. S2D), but not in cholesterol-fed animals (Fig. S1D), we identified an additional peak corresponding to the sterol desmosterol, which accumulated in a stage-specific manner and reached its highest levels during late larval and pupal stages (Figs. 2H, I; S3A). In the final step of the Bloch branch of mammalian cholesterol biosynthesis, the enzyme 24-dehydrocholesterol reductase (DHCR-24) reduces desmosterol to cholesterol. In *C. elegans*, DHCR-24 catalyzes the same biochemical reaction, converting desmosterol derived from plant- and fungal sterols such as stigmasterol and ergosterol into cholesterol to support nematode development^6^. Thus, in sterol auxotrophs, DHCR-24 does not support *de novo* cholesterol synthesis from acetyl-CoA, but instead enables the biochemical conversion of dietary sterols into cholesterol. Building on these findings, the role of DHCR-24 in mediating desmosterol-to-cholesterol conversion has also been demonstrated in a few insect species^23–25^, highlighting the importance of this metabolic strategy across Ecdysozoa. To examine whether a similar mechanism operates in mosquitoes, we searched the *Ae. albopictus* genome for orthologs of human genes involved in cholesterol biosynthesis. Consistent with growth arrest under sterol-free conditions, the *Ae. albopictus* genome displays the sterol auxotrophy signature^6^ characterized by the absence of the first three enzymes in the cholesterol synthesis pathway, FDFT1, SQLE, and LSS (Fig. 3A). The *Ae. albopictus* genome, however, has genes with sequence similarity to several human enzymes participating in cholesterol biosynthesis downstream of FDFT1, SQLE, and LSS activity. Except for *dhcr-24,* the functions of these genes remain uncharacterized in invertebrates^6^. Notably, we identified a clear one-to-one putative ortholog of *dhcr-24*, annotated as LOC109431506 in the AalbF5 assembly of the *Ae. albopictus* genome (Fig. 3A). To examine whether the encoded protein mediates desmosterol-to-cholesterol conversion, we first supplemented larvae with dietary desmosterol. In contrast to vehicle-treated controls, desmosterol robustly supported larval development and survival to adulthood, comparable to cholesterol and plant- and fungal-derived sterols (Figs. 3B; S3A). Progressive accumulation of cholesterol in larvae reared on desmosterol, with levels increasing from larval to pupal and adult stages (Figs. 3C, D; S3B), further supports *in vivo* conversion of dietary desmosterol to cholesterol in *Ae. albopictus*. To test the function of the putative *Ae. albopictus* DHCR-24 orthologue directly, we expressed it in *E. coli* (Figs. 3E; S3C) that neither synthesizes nor modifies sterols and assessed its ability to reduce desmosterol to cholesterol in bacterial lysates. Cholesterol accumulation was detected in lysates from bacteria expressing the candidate *Ae. albopictus* DHCR-24, but not in empty vector controls (Figs. 3F, G; S3D), demonstrating catalytic reduction of desmosterol to cholesterol and biochemically identifying this gene as the *Ae. albopictus* DHCR-24 enzyme.

**Figure 3.**
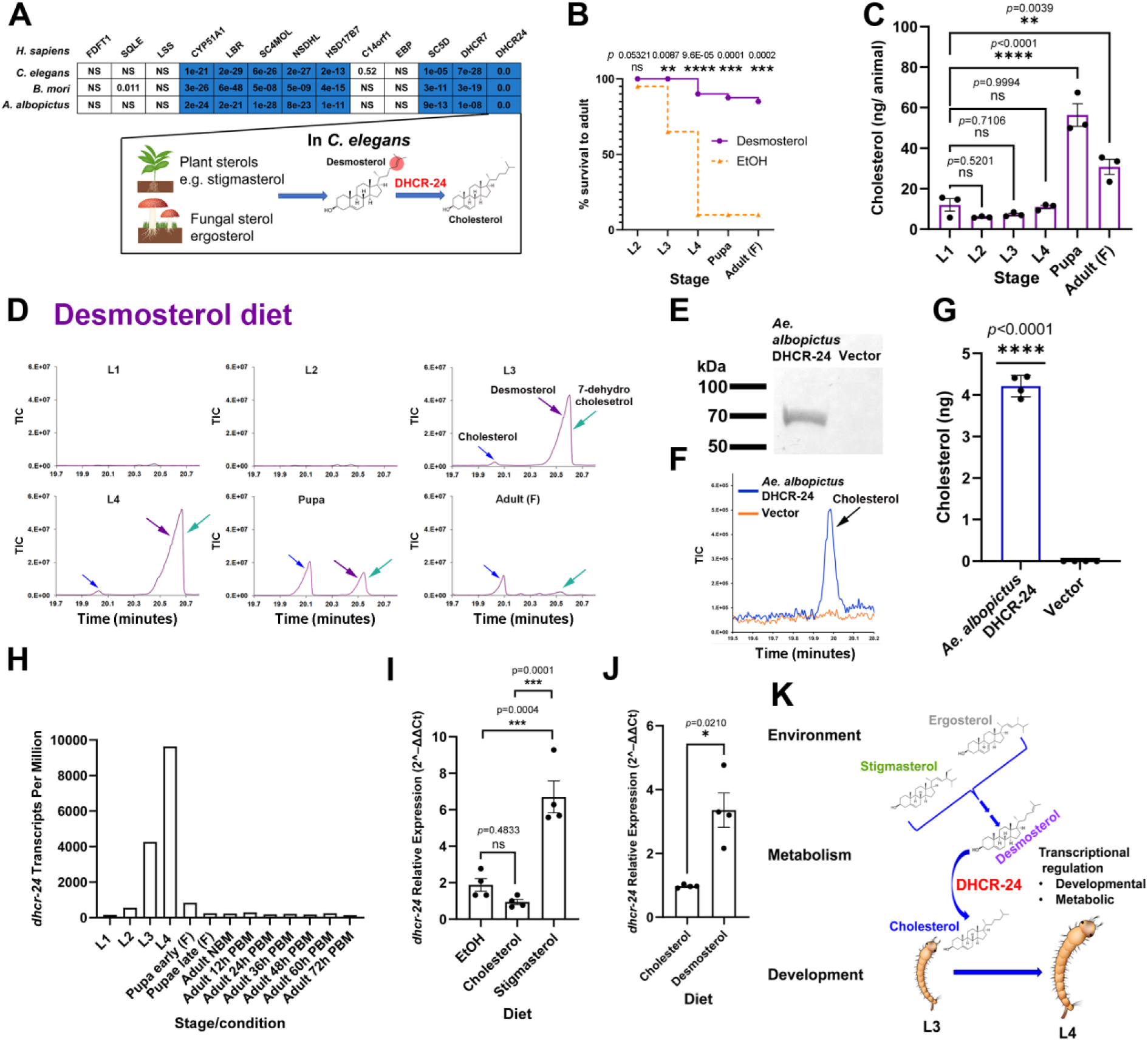
*Ae. albopictus* DHCR-24 facilitates desmosterol-to-cholesterol conversion. **A.** Sequence comparison (BLASTP) of cholesterol biosynthesis enzymes from *Homo sapiens* and related proteins with sequence similarity from *C. elegans, Bombyx mori (B. mori),* and *Ae. albopictus*; NS indicates no significant similarity; squares with E-values ≤ 1 × 10⁻⁶ are highlighted in blue. **B.** Survival at each developmental stage (L2–adult) under desmosterol supplementation (n = 40 individually reared larvae); the EtOH control from Fig. 2A is shown for reference (dotted orange line); Chi-square test. **C.** Cholesterol content per animal across developmental stages in desmosterol-supplemented cultures; n = 3 biological replicates of pooled 20 larvae; one-way ANOVA followed by Dunnett’s post hoc test. **D.** Representative GC–MS sterol chromatograms from 20 larvae reared on a desmosterol-supplemented diet across developmental stages. **E.** Western blot of bacterially expressed *Ae. albopictus* DHCR-24 tagged with a 6×His tag and detected using an anti-His antibody. The uncropped blot is shown in Fig. S3C. **F.** GC–MS chromatograms of sterols extracted from lysates of bacteria expressing *Ae. albopictus* DHCR-24 and empty vector control. The full chromatogram of the sterol-relevant region is depicted in Fig. S3D. **G.** Quantification of cholesterol levels; n = 4 independent cultures per condition; one-sample Student’s *t*-test compared to zero (the level of cholesterol in the empty vector control). **H.** Temporal expression profile of *Ae. albopictus dhcr-24* based on RNA-seq analysis (reference dataset; full profile in Fig. S3E). NBM, no blood meal; PBM, post-blood meal. **I.** *Ae. albopictus dhcr-24* expression measured by qPCR under different dietary conditions; n = 4 independent cultures of pooled 20 L2 larvae harvested 7 days after hatching; one-way ANOVA followed by Tukey’s post hoc test. **J.** *dhcr-24* expression in *Ae. albopictus* measured by qPCR comparing cholesterol and desmosterol diets; n = 4 independent cultures; two-tailed unpaired Welch’s *t*-test. **K.** Model summarizing the proposed role of dietary sterols and DHCR-24-mediated conversion in supporting cholesterol production and developmental progression in *Ae. albopictus* larvae.

Guided by our metabolic analyses showing active desmosterol to cholesterol conversion during larval development, we next asked whether *Ae. albopictus dhcr-24* is developmentally regulated. Examination of a developmental RNA transcriptomic atlas of *Ae. albopictus*^26^ for *dhcr-24* expression revealed a strong, stage-specific upregulation during mid-larval development (L2–L4) (Fig. 3H). This temporal expression pattern coincides with the timing of cholesterol accumulation from the upstream precursor desmosterol in larvae. Notably, *dhcr-24* expression is not altered by blood feeding, consistent with the absence of plant and fungal sterols in blood. The absence of *dhcr-24* upregulation in adult males, adult females, and ovaries (Fig. S3E) further localizes desmosterol-to-cholesterol conversion to the larval stages. To determine whether this stage-specific *dhcr-24* regulation is conserved in related *Aedes* mosquitoes, we compared the expression pattern of the *Ae. albopictus dhcr-24* with the expression of its one-to-one ortholog in *Ae. aegypti*. We observed conserved late-larval upregulation of *dhcr-24* ortholog in *Ae. aegypti* (Fig. S3F), whereas no similar developmental pattern was detected for *C. elegans dhcr-24* (Fig. S3G).

Because *Ae. albopictus* DHCR-24 functions as a sterol-converting enzyme, we reasoned that its expression might be regulated by sterol availability, analogous to substrate- or product-level regulation observed for many metabolic enzymes. To test this hypothesis, we quantified *dhcr-24* transcript levels in larvae grown for five days, reaching the end of the L2 stage, under different sterol-related dietary conditions. No significant differences in *dhcr-24* expression were observed between sterol-depleted and cholesterol-supplemented cultures (Fig. 3I), arguing against product-level regulation or a compensatory response to sterol deprivation. In contrast, supplementation with stigmasterol resulted in significant upregulation of *dhcr-24* expression (Fig. 3I), supporting a substrate-level mode of transcriptional upregulation. Similar upregulation of *dhcr-24* in response to desmosterol (Fig. 3J) suggests direct substrate-dependent positive regulation, linking desmosterol availability to activation of a desmosterol-to-cholesterol conversion mechanism.

Together, these findings are consistent with a model in which *Ae. albopictus* larvae exploit plant- and fungal-derived sterols through a developmentally timed and metabolically regulated DHCR-24–dependent conversion pathway, enabling the production of cholesterol from environmental sterol precursors to sustain late larval growth and development (Fig. 3K).

In light of our finding that the acquisition of dietary sterols is essential for larval development, and given that blood provides a major source of cholesterol, we asked whether cholesterol acquired during blood feeding has a role in *Ae. albopictus* reproduction. To address this possibility, we formulated a sterol-controlled artificial blood meal substitute by depleting sterols from the ingredients of a previously described recipe comprising bovine serum albumin (BSA) as a protein source and phosphatidylcholine (PC) as a lipid source^27^. Thin-layer chromatography (TLC) confirmed efficient removal of sterols from the depleted fraction and selective recovery of cholesterol in the supplemented condition (Fig. S4A). Quantitative sterol analysis further showed that cholesterol levels in sterol-depleted artificial blood were markedly reduced relative to cholesterol-supplemented artificial blood (Fig. 4A, B), validating the effectiveness of sterol depletion. To determine whether cholesterol levels affect the amount of artificial blood consumed, we labeled the artificial blood with fluorescein beads and quantified ingestion in individual females following feeding (Figs. 4C; S4B). The proportion of females that ingested artificial blood, as assessed by fluorescein labeling, did not differ between cholesterol-supplemented and sterol-depleted conditions (Fig. 4D). Quantification of fluorescein levels in fed females further revealed no significant difference in ingestion volume between the two dietary conditions (Fig. 4E), indicating that cholesterol supplementation does not affect feeding propensity or meal size. To determine whether sterol depletion from artificial blood alters cholesterol accumulation in females, we measured sterol levels following artificial blood feeding and found that females fed cholesterol-supplemented artificial blood accumulated substantially higher cholesterol levels than females fed sterol-depleted artificial blood (Fig. 4F), confirming that removal of cholesterol from the artificial blood effectively reduced cholesterol accumulation in females and enabled controlled manipulation of their sterol status. Having established that cholesterol levels in females can be directly controlled through artificial blood feeding, we next asked whether cholesterol availability influences reproductive output (Fig. 4G). Females fed cholesterol-supplemented artificial blood laid significantly more eggs compared with females fed sterol-depleted artificial blood (Figs. 4H; S4C–E), although egg production remained lower than that observed following feeding on blood. Temporal analysis indicates that differences between cholesterol-supplemented and sterol-depleted artificial blood are concentrated in the first two days after feeding, when most eggs are laid (Fig. S4D). These results identify cholesterol as a limiting reproductive nutrient acquired through blood feeding. Cholesterol availability in the adult diet is therefore a key determinant of reproductive capacity in *Ae. albopictus*.

**Figure 4.**
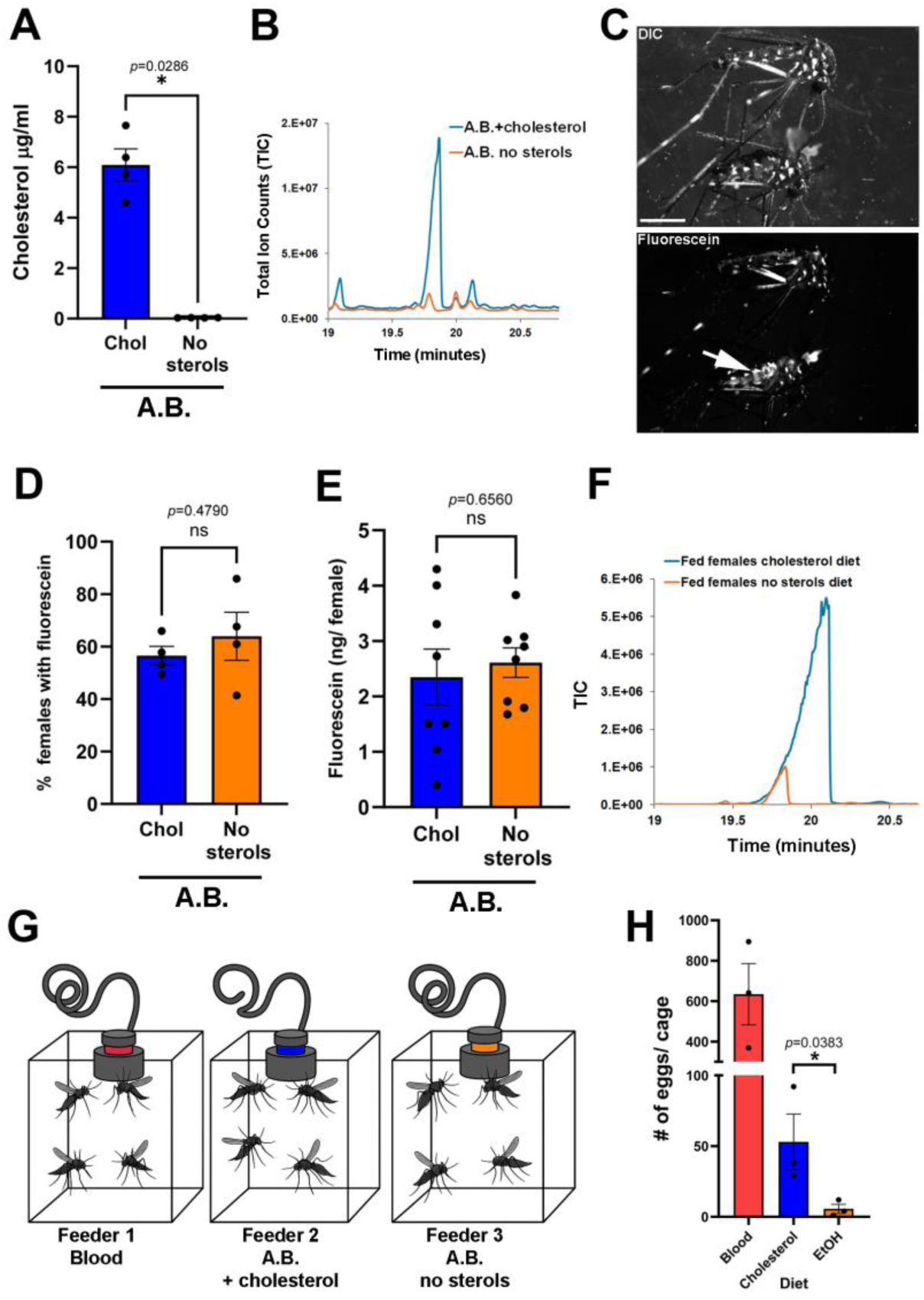
Dietary cholesterol depletion reduces egg laying. **A.** Cholesterol concentration in artificial blood under cholesterol-supplemented or sterol-depleted conditions (n = 4 independent samples; unpaired two-tailed Mann–Whitney test). **B.** Representative GC–MS chromatograms of cholesterol-supplemented or sterol-depleted artificial blood. **C.** Representative DIC and fluorescence images of fed females; white arrow indicates fluorescence signal in the female intestine; scale bar, 1 mm. **D.** Percentage of fluorescein-positive females after feeding on cholesterol-supplemented or sterol-depleted artificial blood; n = 4 independent experiments per condition; unpaired two-tailed Student’s *t*-test. **E.** Fluorescein levels in females fed fluorescein-labeled cholesterol-supplemented or sterol-depleted artificial blood; n = 8 independent females per condition; unpaired two-tailed Student’s t-test. **F.** Representative GC–MS chromatograms from pooled samples of 20 females per condition. **G.** Schematic of the artificial blood-feeding system. **H.** Number of eggs laid per cage under blood-fed, cholesterol-supplemented artificial blood, or sterol-depleted artificial blood conditions; unpaired one-tailed Student’s *t*-test.

Integrated with our developmental analyses, these findings unveil a stage-specific sterol metabolic cycle in mosquitoes, in which egg cholesterol, plausibly of maternal origin, sustains early larval development, later larval stages convert environmental sterols to cholesterol, and adult females acquire host-derived cholesterol to drive egg laying.

## Discussion

Using *Ae. albopictus* as a model, we developed an experimentally integrated system to study sterol metabolism across the mosquito life cycle. Our results identify a developmentally coordinated sterol conversion axis that regulates cholesterol availability according to stage-specific physiological demands. Cholesterol in eggs supports early larval development, whereas later larval stages utilize plant- and fungal-derived sterols that are metabolically converted into cholesterol by the activity of DHCR-24. This conversion is temporally regulated to coincide with the developmental window during which dietary sterols are required to sustain larval growth. In adult females, cholesterol derived from the blood meal is essential for efficient egg laying. Together, these findings show that mosquitoes flexibly exploit distinct environmental sterol sources across life stages, linking stage-specific physiology with the metabolic logic of cholesterol acquisition and utilization.

We found that cholesterol levels vary markedly in eggs from different mosquito species, with no clear correlation with oviposition strategy (egg rafts versus individual eggs) or with the growth conditions of the females (wild versus laboratory-reared mosquitoes). Notably, some mosquito eggs contain cholesterol at a level approximately 0.4% of egg mass, comparable to the typical 0.4–0.5% reported for chicken eggs^28^. We propose that sterol auxotrophy, together with the need to provision eggs with substantial amounts of cholesterol for early larval development, may have contributed to the evolutionary emergence of host blood feeding as a dietary source of egg cholesterol. Although mammalian blood sterols are composed largely of cholesteryl esters (CEs), which constitute approximately 70% of total sterols^29^, mosquito eggs are enriched in free cholesterol, and CEs were not detected. This observation suggests intestinal hydrolysis of cholesteryl esters (CEs) prior to cholesterol deposition in the egg, consistent with a mechanism reported in *Helicoverpa armigera*^30^. Our results show that egg cholesterol supports early larval development and thus can enable growth in habitats that are initially sterol-limited until environmental sterols accumulate through the progressive buildup of organic material^31^. Differences in egg cholesterol levels may therefore influence colonization success. Habitats that are initially sterol-poor, including tires and plastic containers, are commonly colonized by *Ae. albopictus* larvae^32^. Our findings suggest that egg cholesterol reserves may facilitate early establishment in such environments, although high egg cholesterol levels are not unique to Ae. albopictus (Fig. 1A) and likely act together with additional ecological and physiological adaptations.

In addition to its established role as a precursor for the molting hormone ecdysone, cholesterol has been implicated in molting-independent functions in *C. elegans*^33^ and *Drosophila*^16^. Consistent with this notion, no obvious molting defects were observed under cholesterol depletion in *Ae. albopictus*, suggesting that cholesterol is required beyond ecdysone production during mosquito development. Plant and fungal sterols are likely relevant sterol sources in natural habitats, as not all such environments contain cholesterol^2^. Notably, cholesterol levels were lower in stigmasterol-fed larvae compared to cholesterol-fed larvae, consistent with metabolic conversion representing a possible bottleneck for cholesterol availability. Plant sterols such as beta-sitosterol have been proposed to act as larvicidal compounds at a concentration of 27 µM^34^, indicating that concentration is an important determinant of their biological activity. Our results suggest that in natural environments, plant sterols such as stigmasterol may support, rather than inhibit, mosquito development.

The role of DHCR-24 as an enzyme facilitating the conversion of desmosterol to cholesterol has been demonstrated in a few species of insects^23–25^. Our results demonstrate that in *Aedes* mosquitoes, this pathway is regulated developmentally and metabolically, potentially coordinating the level of dietary sterol utilization with their availability. Based on studies in *C. elegans* and Bombyx *mori* showing that *dhcr-24* is expressed in the intestine^6,35^, the primary site of sterol uptake, sterol conversion in *Aedes* mosquitoes is also likely to occur in the intestine. Notably, DHCR-24 expression was not upregulated in *Ae*. *albopictus* adult males, despite their consumption of plant-derived sterols^36^ through nectar feeding, suggesting that desmosterol-to-cholesterol conversion is restricted to late larval stages. The upregulation of *dhcr-24* transcription by stigmasterol and desmosterol but not by cholesterol or its depletion suggests a sterol-encoded mechanism of regulated transcription. The transcription factors involved, and the mechanisms by which specific sterols activate this response, remain to be elucidated, as sterol-specific regulation of transcription in invertebrates has remained largely unexplored.

Intriguingly, unlike *Aedes* mosquitoes and other *Culicines*, we did not identify an obvious ortholog of human or *Aedes* DHCR-24 in available *Anophelinae* genomes. This suggests that, as reported in some insects^7^, *Anophelinae* may utilize plant and fungal sterols directly rather than converting them to cholesterol. Alternatively, these mosquitoes might possess a DHCR-24-independent sterol conversion pathway or rely exclusively on dietary cholesterol. Our findings in *Aedes* mosquitoes provide a molecular and metabolic basis for understanding divergent and convergent mechanisms of sterol utilization across *Culex*, *Culiseta*, and *Anopheles* mosquitoes.

By extending sterol analysis to adult feeding and reproduction, we show that cholesterol availability in the adult diet directly constrains egg laying. This finding provides functional support for an earlier study in *Ae. aegypti*^27^ in which dietary supplementation of low-density lipoprotein or a phospholipid–cholesterol mixture supported egg laying in adult females. Our findings of stage-specific acquisition of distinct dietary sterols across the mosquito life cycle provide a functional context for previous reports describing the activity and regulation of sterol- and oxysterol-binding carrier proteins in mosquitoes. For example, upregulation of the sterol carrier protein SCP-2 in the intestine of L3 and L4 *Ae. aegypti* larvae have been reported^37^, suggesting active dietary sterol acquisition at these stages, consistent with our findings. Blood meals induce upregulation of genes related to sterol transport, including increased expression of several oxysterol-binding proteins in the ovary of *Ae. aegypti*^38^ and SCP proteins in the midgut, fat body, and ovary of *Anopheles stephensi*^39^. In addition, multiple sterol transporters are upregulated in a blood-meal-dependent manner in the midgut of *Ae. aegypti* and *Anopheles gambiae*^40^, collectively suggesting enhanced and regulated transport of blood-derived sterols from the midgut to the ovaries. Our study reveals the biological significance of this regulated sterol transport in enabling efficient egg laying.

While the role of nutrition-derived proteins in mosquito development and reproduction has been reported^41,42^, our study establishes stage-specific sterol acquisition and utilization as an additional essential nutritional axis in mosquitoes. Defining sterol metabolic pathways in mosquitoes is increasingly recognized as central to mosquito vectorial capacity and, consequently, to human health. For example, the *Plasmodium* life cycle is sensitive to the level of cellular cholesterol of its mosquito host^39^. Similar crosstalk between mosquitoes, their viruses, and cholesterol has been reported in the case of dengue^43^ and Zika viruses^44,45^. In addition, targeting of insect cholesterol metabolism, for example, via the manipulation of SCP proteins, may represent a promising strategy for larvicide development^46,47^. Together, our findings advance the molecular and metabolic understanding of sterol metabolism across the life cycle of *Aedes* mosquitoes, identifying sterol acquisition and conversion as physiologically constrained processes that represent potential vulnerabilities for disrupting mosquito development and reproduction in ecological and disease-control contexts.

## Acknowledgments

We thank Jonathan D. Bohbot and Evyatar Sar-Shalom (The Hebrew University of Jerusalem) for providing *Ae. aegypti* eggs, and Ido Tsurim (Achva Academic College) for providing mosquito feeders. We thank Diptera.ai for supplying Ae. albopictus eggs and for conducting the egg-laying assays, and Matan Ben-Ari (Oranim College of Education) for establishing egg sampling sites at the Oranim botanical garden. We are grateful to Benjamin Podbilewicz (Technion – Israel Institute of Technology) for critically reading the manuscript. This work was supported by the Israel Science Foundation (ISF) grant 779/21 to Amir Sapir and by ISF grant 1166/22 to Alon Silberbush and Yoram Gerchman.

## Author contribution

S.C., B.T., Y.G., A.Si. (Silberbush), and A.Sa. (Sapir) designed the research; B.T., D.S.Y.Z., P.A.P., and A.Sa. performed DHCR-24 expression analysis; S.C., B.T., Y.G., A.Si., and A.Sa. conducted the mosquito experiments and analyzed the data; A.Sa. wrote the paper.

## Methods

### Cholesterol quantification in mosquito eggs

Eggs of *Ae. albopictus* were obtained from Diptera Ltd. (Jerusalem, Israel) and from the Papathanos laboratory (which also provided *Anopheles gambiae* eggs). *Ae. aegypti* London strain eggs were obtained from the Bohbot laboratory at the Hebrew University. Egg rafts were collected from natural ponds and water containers at Oranim Academic College of Education (Kiryat Tivon, Israel). Individual rafts were imaged, and a small subset of eggs (∼20) was removed for hatching, while the remaining eggs were transferred to microcentrifuge tubes and stored at −80 °C until analysis. Larvae were reared to the L2 stage, and species identity was determined morphologically. The number of eggs per raft used for GC–MS analysis was determined by subtracting the number of eggs removed for hatching from the total egg count obtained from raft images.

### Egg hatching and larval growth conditions

Eggs were hatched overnight in double-distilled water (DDW) under controlled laboratory conditions at 28 °C with a 12:12 hours (h) light: dark cycle. Larvae were collected exactly 24 h post-hatching to ensure developmental synchrony and were used for all subsequent experiments.

### Sterol supplementation

To assess the effects of dietary sterols on larval survival, development, and gene expression, sterols were supplemented either at hatching or at defined developmental time points, depending on the experimental design. The following sterols were used: cholesterol (Cat# 26732, Sigma-Aldrich, purity 99%), stigmasterol (Cat# sc-281156, Santa Cruz Biotechnology, purity ≥ 95%), ergosterol (Cat# 45480, Sigma-Aldrich, purity ≥ 95%), and desmosterol (Cat# 14943, Cayman Chemical, purity 98%). Sterols were dissolved in ethanol (Cat# 05250501, Biolabs-Biology) and added to the rearing water at a final concentration of 13 µM. Control groups received ethanol alone, with solvent concentrations matched across all treatments.

### Preparation of chemically defined media for bacterial growth

All solutions were prepared using HPLC-grade double-distilled water (Cat# 002321060200, Biolabs-Biology). M9 salt solution, which serves as the basis of minimal essential medium (MEM), was prepared according to a previously published protocol^1^. Briefly, 42 mM Na₂HPO₄, 22 mM KH₂PO₄, 86 mM NaCl, and 19 mM NH₄Cl were dissolved and supplemented with 1 mM MgSO₄ and 0.1 mM CaCl₂. The pH was adjusted to 7.2 using NaOH or KOH prior to sterilization by autoclaving for 20 minutes (min). Solutions were stored at room temperature until use. M9-based minimal medium was prepared by diluting M9 salts five times in water and supplementing with glucose (0.4% w/v), MgSO₄ (1 mM), and CaCl₂ (0.1 mM). Media were prepared under sterile conditions and used as specified in the experimental procedures.

### Bacterial growth and larval feeding

Escherichia coli strain MG1655 was cultured in MEM for 24 h at 37 °C with shaking. Cultures were harvested, and bacterial pellets were washed twice with DDW before being adjusted to a final OD₆₀₀ of 0.05. Bacteria were supplied to all experimental conditions at this density and replenished every 48 h throughout larval development.

### Population and individual assays

For population-based survival assays, larvae were randomly distributed into plastic cups containing 400 mL of DDW supplemented with bacterial feed (30 larvae per cup). For individual-level developmental analyses, sterol-timing experiments, and molecular assays, single larvae were transferred into glass vials containing 5–7 mL DDW with bacterial feed. All experiments were conducted under controlled environmental conditions at 28 °C with a 12:12 h light: dark cycle. Water levels were monitored daily and replenished with DDW as needed.

### Larval survival and developmental monitoring

Larval survival and development were monitored daily. Dead larvae were removed and recorded. Developmental progression was assessed visually, and transitions between larval instars were determined by the presence of exuviae (molted cuticles) in the water. Pupae were transferred individually to test tubes to prevent cross-contamination and enable accurate sex determination. Adult emergence was recorded, and sex was determined upon emergence. This monitoring procedure was applied to both continuous sterol treatments and time-specific sterol supplementation assays.

### Periodic sterol supplementation

To examine the effect of supplementation timing on larval development, individual larvae were maintained in glass vials containing 7 mL of DDW and bacterial feed. *E. coli* MG1655 was replenished every 48 h at OD₆₀₀ = 0.05. Larvae received a single supplementation of either cholesterol or stigmasterol at defined developmental time points (day 1, 4, 8, 12, or 16 post-hatching). Larvae were monitored daily until pupation and adult emergence.

### Effect of sterol exposure duration on larval development and survival

To assess the impact of sterol exposure duration, newly hatched larvae were distributed into plastic cups (10 larvae per cup) containing 400 mL of DDW and bacterial feed. Twenty-four hours post-hatching, cholesterol or stigmasterol was added to each cup at the designated concentration. After four days of exposure, larvae from six cups per sterol treatment were transferred to fresh cups containing DDW and bacterial feed. In three cups per treatment, sterol supplementation was continued at the same concentration, whereas larvae in the remaining three cups were transferred to sterol-free DDW supplemented with bacterial feed and ethanol as a solvent control. This transfer procedure was repeated at four-day intervals until all individuals either died or reached adulthood. At the conclusion of the experiment, survival was quantified for each treatment group. Environmental conditions were identical to those described above.

### Larval imaging

Larvae at defined developmental stages or experimental time points were collected and fixed in 100% ethanol to preserve morphology. Fixed specimens were washed twice with distilled water to remove residual ethanol and minimize potential distortion. Individual larvae were transferred to 55 mm transparent plastic plates and positioned to prevent overlap during imaging. Imaging was performed using a Nikon SMZ18 stereomicroscope using the NIS-Elements software. All imaging parameters and acquisition settings were kept constant across developmental stages and experimental replicates to enable direct comparison of morphological features.

### Gene expression analysis by quantitative PCR (qPCR)

For gene expression analyses, 30 larvae per sample were reared under defined sterol conditions as described above and sampled on day 5 of development. Total RNA was extracted using TRIzol reagent (Cat# 1141831, Biolabs-Biology) in combination with the DirectZol RNA kit (Cat# R2072, Zymo Research). Briefly, 200 µL TRIzol was added to each sample, and homogenization was performed using a motorized pestle for 2 min with samples kept on ice. An additional 400 µL TRIzol was added, followed by incubation for 5 min at room temperature. Subsequently, 120 µL chloroform was added, samples were vortexed for 15 s and incubated for 15 min at room temperature before centrifugation at 14,000 × g for 15 min at 4 °C. The aqueous phase was transferred to an RNase-free tube, mixed with an equal volume of 95–100% ethanol, and loaded onto a Zymo-Spin™ IIICG column. RNA purification was completed according to the DirectZol protocol, including on-column DNase treatment. RNA concentration was quantified using a Qubit® 3.0 Fluorometer (Thermo Fisher Scientific). One microgram of total RNA was used for complementary DNA (cDNA) synthesis using the qScript cDNA Synthesis Kit (Cat# 95047-100, QuantaBIO). Quantitative PCR reactions were performed using PowerTrack™ SYBR Green Master Mix (Cat# A46109, Applied Biosystems) on a QuantStudio™ 1 Real-Time PCR System (Thermo Fisher Scientific). Primers were designed using the National Center for Biotechnology Information (NCBI) Primer-BLAST tool or selected based on published sequences². All primers were synthesized by Integrated DNA Technologies (IDT) and are listed in Table 1 together with their melting temperatures and amplicon sizes. Each 10 µL reaction contained 5 µL PowerTrack™ SYBR Green Master Mix, 0.2 µM of each primer, 4 ng cDNA, and RNase-free water to volume. PCR cycling conditions were as follows: 95 °C for 20 seconds (s), followed by 40 cycles of 95 °C for 1 second and 60 °C for 20 seconds (annealing/extension). Melt-curve analysis was performed by heating to 95 °C for 1 second, cooling to 60 °C for 20 seconds, reheating to 95 °C for 1 second, and cooling again to 60 °C for 20 seconds to verify amplification specificity. Amplification efficiency (E) and correlation coefficients (R²) were determined for each primer pair using fivefold serial dilutions of 50 ng cDNA (1:5, 1:25, 1:125, 1:625, and 1:3,125) to generate standard curves. Amplification efficiency was calculated using the formula: E = (10^(–1/slope) − 1) × 100. Acceptable primer efficiency ranged between 95% and 115%. Gene expression levels were normalized to the ELF-1g transcript^2^ and calculated using the ΔΔCt method. Results represent the mean of at least five biological replicates, and error bars indicate standard errors (SE). No-template controls were included in all runs to exclude contamination.

**Table 1.**
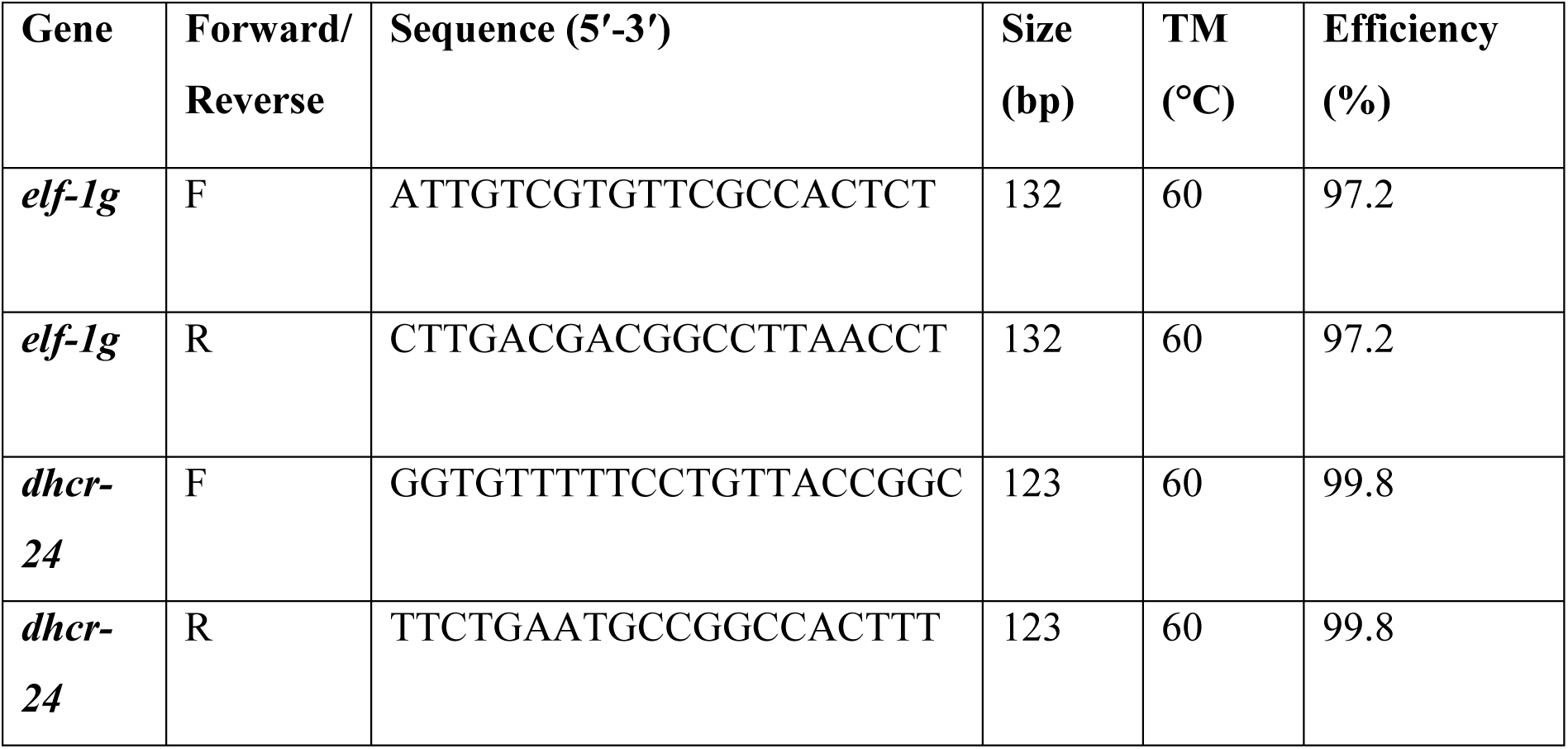
Primers used in this study.

### Preparation of mosquito larvae for gas chromatography sterol profiling

To assess sterol composition by gas chromatography-mass spectrometry (GC-MS), *A. albopictus* eggs were hatched as described above. Individual larvae were transferred into glass vials containing 7 mL of double-distilled water (DDW) supplemented with bacterial feed, and sterol treatments were applied according to the protocol described above. Larval development was monitored daily by scoring exuviae to determine progression through each instar, from hatching to pupation and adult emergence, of which only adult female were collected for GC analysis. For each developmental stage (egg, instar I–IV, pupa, and adult females), 30 individuals were pooled per sample. Each condition was analyzed in three biological replicates, with each replicate consisting of 30 individuals derived from the same treatment group and originating from a single rearing vial. Replicates were obtained from the same egg batch collected from a single rearing cage to minimize genetic variability within and between replicates. Specimens were briefly rinsed with DDW to remove external contaminants and stored at −80 °C until processing. All samples were processed under identical conditions to minimize technical variability.

### Quantification of sterols

Sterol quantification was performed based on a previously described protocol¹ with minor modifications. Briefly, 100 µL H₂O was added to each sample, followed by homogenization using a motorized pestle for 2 min. Subsequently, 500 µL analytical-grade ethanol containing 5 µg of the internal standard campesterol (Avanti Polar Lipids, Cat# 700122) was added. Fifty microliters of freshly prepared 10 N NaOH in H₂O was then added, tubes were sealed with parafilm, and samples were incubated at 70 °C for 1 h, followed by 3 min at 43 °C. Phase extraction was performed by adding 250 µL H₂O and 500 µL hexane (BioLabs, Cat# 08290601). Tubes were sealed, vortexed for 1 min, and centrifuged at 4,000 × g for 5 min. The upper organic phase (∼500 µL) was collected, and extraction was repeated twice. The combined hexane fractions (∼1.5 mL) were washed with 500 µL H₂O and centrifuged as described above. The organic phase was transferred to GC glass vials, dried under nitrogen gas, and derivatized with 50 µL N, O-bis(trimethylsilyl)trifluoroacetamide (BSTFA; Sigma-Aldrich, Cat# 155195) at 80 °C for 30 min. Subsequently, 150 µL hexane was added to each vial. Five microliters of each sample were injected into an Agilent 7890A gas chromatograph (Agilent Technologies) equipped with an Rxi-5Sil MS capillary column (30 m length, 0.25 mm ID, 0.25 µm film thickness; Restek) and coupled to an Agilent 5975C single-quadrupole mass spectrometer. The oven temperature program was as follows: initial temperature 70 °C; ramped at 20 °C/min to 250 °C; then at 10 °C/min to 320 °C; held at 320 °C for 10 min. Injector temperature was set to 280 °C. Helium was used as a carrier gas at a constant flow of 1 mL/min. Splitless injection was used with a purge time of 1 min. The transfer line temperature was set at 320 °C. Mass spectrometry conditions were: electron impact ionization (70 eV), ion source temperature 230 °C, quadrupole temperature 150 °C, and scan range 40–550 m/z. Sterols were identified based on retention time and comparison of fragmentation spectra with commercial standards and reference spectra from the NIST 08 library. Quantification was performed by calculating the ratio of the sterol peak area to the peak area of the internal standard and applying this ratio to the linear regression equation derived from the corresponding calibration curve (Fig. S2B). Calibration curves were generated using purified commercial sterol standards dissolved in ethanol at concentrations ranging from 3.125 to 50 µg/mL, each containing 5 µg internal standard. Standards were dried, derivatized as described above, and analyzed by GC–MS. Linear regression equations were used to determine sterol concentrations in experimental samples.

### Assay for *Ae. albopictus* DHCR-24 activity in bacteria

The coding region of *Ae. albopictus* DHCR-24 was cloned into pET28a by gene synthesis (Twist Bioscience), and the construct was verified by sequencing. pET28a-DHCR-24 and empty vector control were transformed into BL21 (DE3) bacteria and plated on LB agar containing 50 μg/ml kanamycin. For each strain, a single colony was inoculated into 2 ml MEM buffer supplemented with kanamycin and grown overnight at 37 °C with shaking. The following day, cultures were diluted 1:20 in MEM buffer containing kanamycin and grown for 4 h at 37 °C with shaking until the optical density at 600 nm reached 0.5. Protein expression was induced by adding isopropyl β-D-1-thiogalactopyranoside (IPTG; BioLabs, Cat. #1010) to a final concentration of 1 mM, and cultures were incubated for an additional 4 h at 37 °C in the dark. Cells were harvested by centrifugation at 4,000 g for 20 min at 4 °C, and pellets were resuspended in 500 μl recovery buffer (MEM supplemented with SIGMAFAST protease inhibitor cocktail, EDTA-free; Sigma-Aldrich, Cat. #S8830). Samples were transferred to microcentrifuge tubes and lysed by sonication (Ultrasonic Probe Sonicator, Sonics, Cat. #VCX130) at 20% amplitude on ice, using six cycles of 10 seconds sonication followed by 15 seconds recovery. Desmosterol (5 μl of a 13 mM stock) was added to a final concentration of 130 µM, and samples were incubated at 30 °C in the dark for 12 hours, then stored at −20 °C. For downstream analyses, 2 μl was used for protein detection by standard western blotting, and 400 μl was subjected to GC–MS analysis.

### Preparation of artificial blood food

Artificial blood food was prepared based on a previously described protocol^3^ with minor modifications. Prior to micelle preparation, phosphatidylcholine was purified to remove contaminating sterols according to a published method^4^. Briefly, 5 grams of phosphatidylcholine was dissolved in 30 mL of chloroform. A Bond Elut aminopropyl (NH₂) cartridge column (10 g; Cat# 14256036, Agilent) was equilibrated by rinsing twice with 20 mL hexane. The phosphatidylcholine solution was loaded onto the column and allowed to pass through. The column was subsequently washed with 40 mL chloroform containing 5% (v/v) 2-propanol, followed by 40 mL chloroform containing 7.5% (v/v) 2-propanol. Phosphatidylcholine was eluted with 40 mL of methanol. A 50 µL aliquot was retained for thin-layer chromatography (TLC) analysis to assess sterol removal, and the remaining eluate was dried under a stream of nitrogen.

For TLC verification, methanol-solubilized lipids were applied dropwise onto silica gel 60 TLC plates (Merck; aluminum-backed). Plates were first developed in chloroform:methanol:water (60:30:5, v/v/v) until the solvent front reached mid-height. After air drying, plates were redeveloped in hexane: diethyl ether: acetic acid (80:20:1.5, v/v/v) up to approximately 5 mm from the top. Lipids were visualized by immersion in 10% (v/v) H₂SO₄ followed by heating at 90 °C until bands appeared. Purity was assessed by comparison to unpurified phosphatidylcholine and a cholesterol standard loaded on the same plate. To prepare cholesterol-containing micelles, 300 mg cholesterol and 2100 mg purified phosphatidylcholine were dissolved in 30 mL chloroform, thoroughly mixed, and dried under nitrogen in a round-bottom flask to form a lipid film. The film was resuspended in 75 mL 2× Tyrode’s buffer (16 g NaCl, 0.4 g CaCl₂, 0.2 g MgCl₂, 0.4 g KCl, 0.1 g NaH₂PO₄, and 2 g glucose per liter; pH 7.4) by vigorous vortexing, followed by 30 min sonication until complete emulsification. Control phospholipid micelles lacking cholesterol were prepared using the same procedure, omitting cholesterol. To the lipid suspension, 75 mL of 20% bovine serum albumin (BSA) containing 200 µg/mL fluorescein was added. Fluorescein served as a tracer to estimate the amount of food consumed.

### Assessment of feeding activity and cholesterol uptake in adult females

To assess feeding activity and cholesterol uptake, approximately 100 females and 10 males were introduced into cubic cages (20 × 10 × 10 cm). Adults were deprived of food for 48 h prior to the experiment, with continuous access to water. Mosquitoes were then provided with artificial blood via an overhead feeder for 24 h. The experiment was conducted in three independent experimental blocks, each including treatments #2 and #3 described in Table 2. Forty-eight hours after feeding, cages were placed at −20 °C for 10 min to immobilize the mosquitoes. Female mosquitoes were collected and counted to determine the total number of females and the proportion that had consumed the artificial diet. Feeding status was assessed using fluorescein as an ingestion marker. Fed females were identified by intestinal fluorescence under a binocular dissecting microscope (Nikon SMZ8) equipped with a fluorescence filter set. In addition, 20 fed females per treatment were collected for subsequent GC–MS analysis to quantify cholesterol levels.

**Table 2.** The different diets used for feeding *A. albopictus* adults.

| **Diet** | **ATP** | **Sucrose** | **BSA**<br>**10%** | **Cholesterol**<br>**2mg/ml** | **PC**<br>**70mg/ml** | **Tyrode's**<br>**buffer** | **Fluorescein** |
| --- | --- | --- | --- | --- | --- | --- | --- |
| **1** | blood |  |  |  |  |  |  |
| **2** | + | + | + | - | + | + | + |
| **3** | + | + | + | + | + | + | + |

### Estimation of artificial blood meal consumption using fluorescein

To estimate the volume of artificial blood consumed, ten fed females selected based on intestinal fluorescence were transferred individually to Eppendorf tubes containing 100 µL H₂O and homogenized using a motorized pestle for 2 min. Five microliters of each homogenate were transferred to a black 96-well plate. To each well, 45 µL H₂O and 50 µL 50 mM phosphate buffer (pH 7.4) were added. Fluorescence was measured from the bottom using an EZ Read 400 plate reader (Biochrom Ltd., UK) with excitation at 485/20 nm, emission at 528/20 nm, and gain set to 35. The volume of artificial blood consumed was calculated using the linear regression equation derived from a calibration curve. For the construction of the calibration curve, defined volumes of artificial blood (0.039–10 µL) were added to wells, adjusted to 50 µL with distilled water, and mixed with 50 µL 50 mM phosphate buffer (pH 7.4). Fluorescence was measured under identical instrument settings, and fluorescence values were plotted against known volumes to generate the standard curve used for quantification.

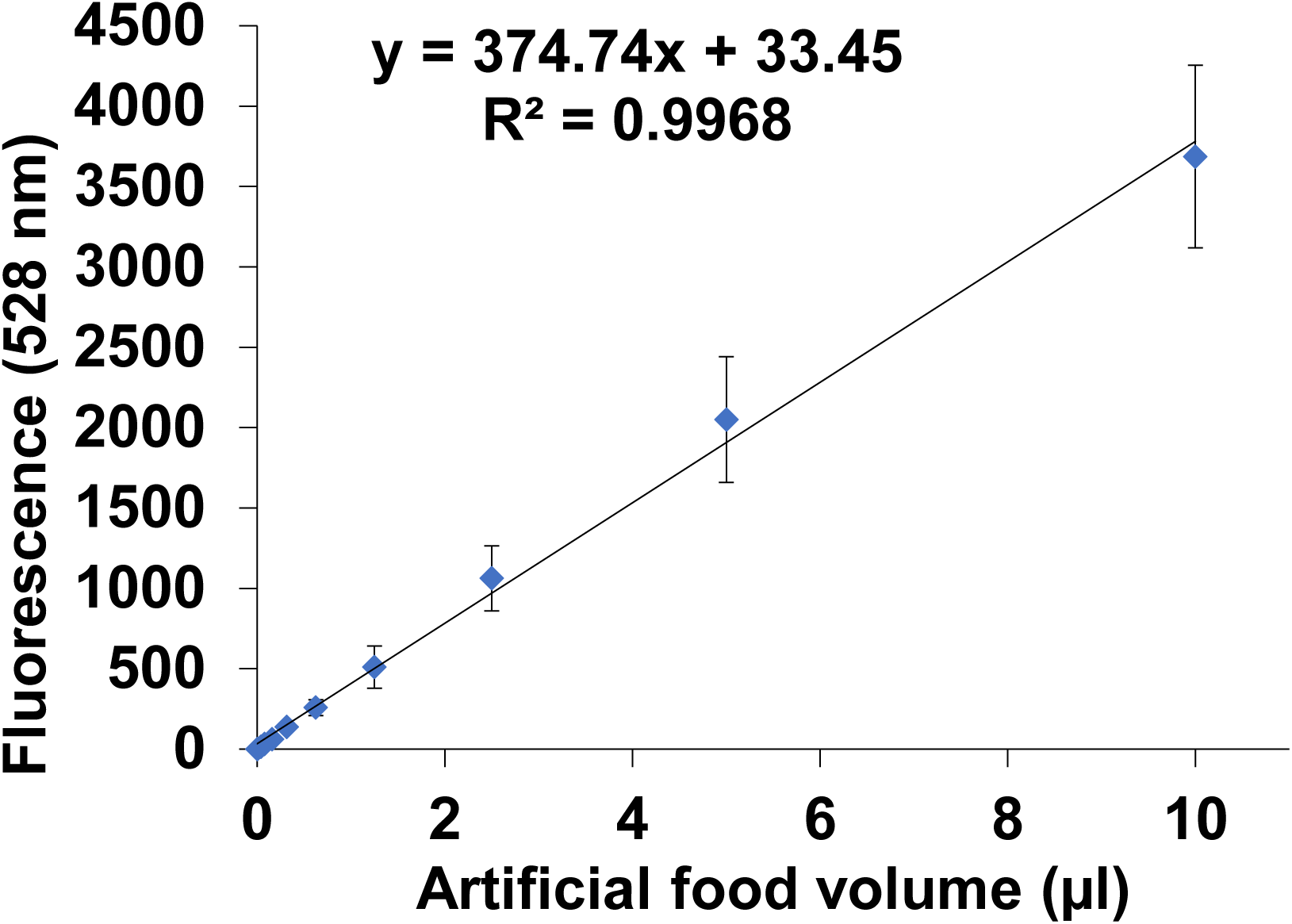

### Effect of dietary cholesterol on egg laying in adult females

Adult *A. albopictus* were maintained in cages (20 × 20 × 20 cm), each containing 500 females and 50 males. Prior to feeding, mosquitoes were deprived of food for 48 h with access to water only. Adults were then provided with the experimental diet via an overhead feeder for 24 h. The experiment was conducted in three independent blocks, each including all dietary treatments as specified in Table 2, with added ATP (1µg/ml) and glucose (10 g/ liter) to each diet. Following feeding, an oviposition container containing 300 mL of DDW and lined with oviposition paper was placed in each cage. Oviposition papers were collected at four 48-h intervals, and eggs were counted. All egg production experiments were performed at the Diptera Company laboratory (Jerusalem, Israel).

### Statistical analysis and graphs

Statistical analyses were performed using Prism (GraphPad Software) or SPSS (IBM). Data visualization was conducted using Prism, figures were assembled in Adobe Photoshop (Adobe Systems), and the concluding model was illustrated using Adobe Illustrator.

