## Supplementary data for "Cholesterol links blood feeding to mosquito development and reproduction"

**Supplementary figures**

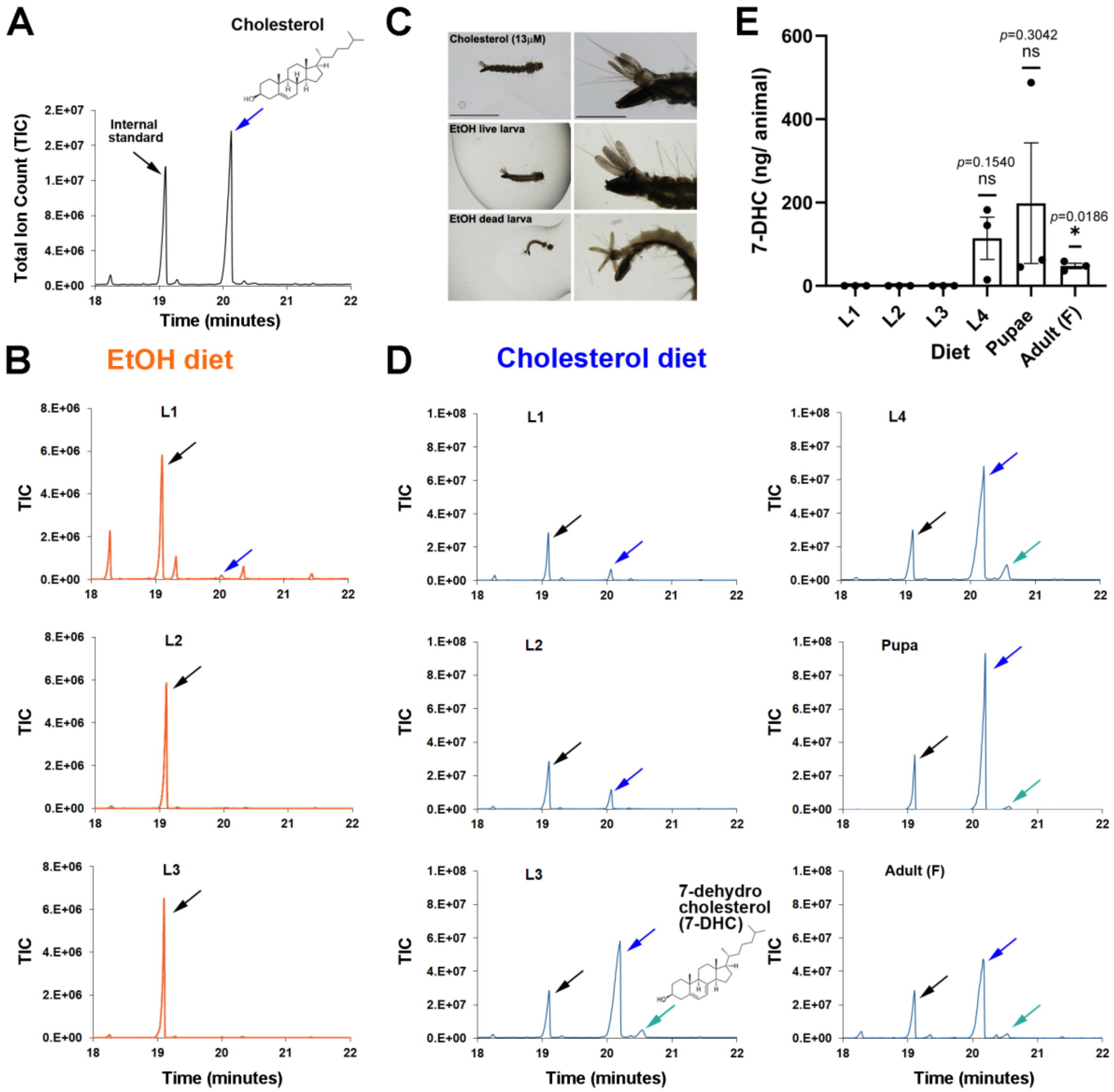


**Figure S1. Cholesterol and 7-dehydrocholesterol identification. A.** Representative GC–MS chromatograms of sterols extracted from *Ae. albopictus* eggs showing cholesterol and coprostanol that was used as an internal standard (I.S.) for sterol identification and normalization. Hereafter, the sterol region (18–22 minutes) of the chromatograms is shown. **B.** Representative GC–MS chromatograms of EtOH-supplemented cultures corresponding to Fig. 1D. **C.** Representative images of live and dead larvae from sterol-defined cultures. Scale bars, 5 mm for the right panels and 1 mm for the left panels. **D.** Representative GC–MS chromatograms of cholesterol-supplemented cultures corresponding to Fig. 1J. **E.** Quantification of 7-dehydrocholesterol in the samples shown in Fig. 1J. Statistical significance was assessed using a one-sample Student's *t*-test against zero, as 7-DHC was undetectable at L1.

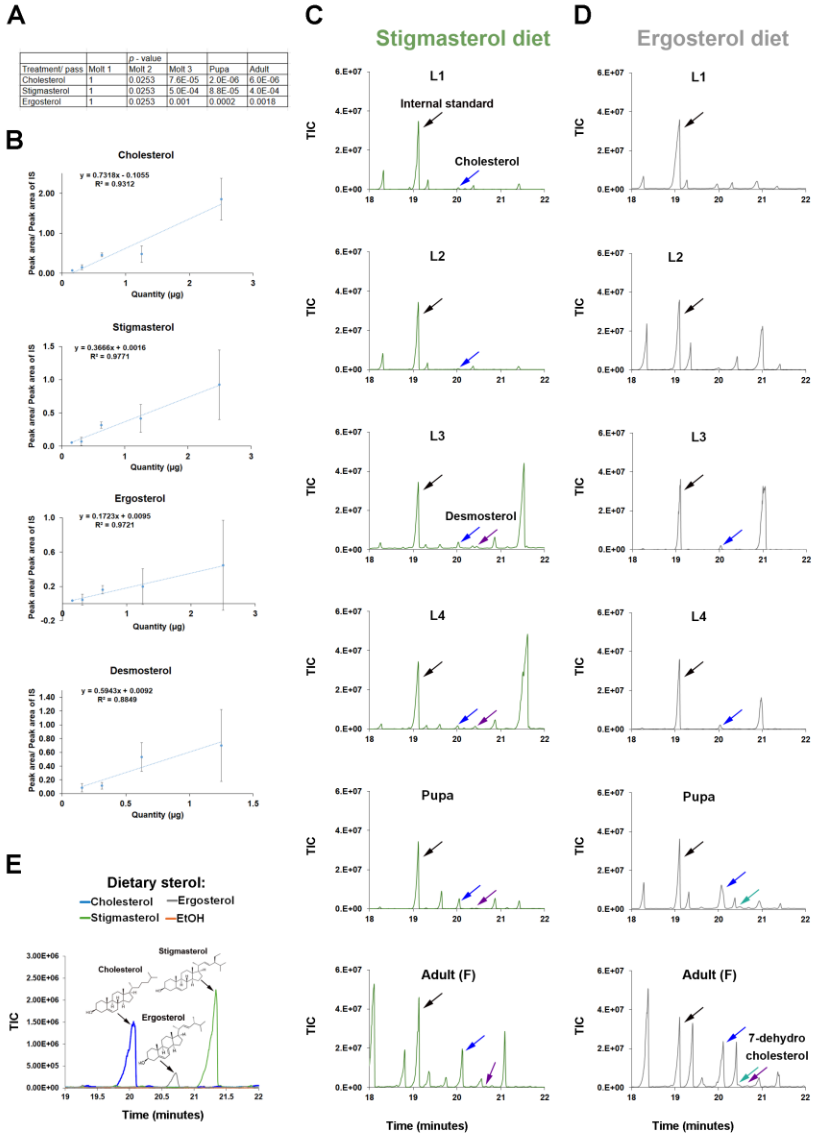


**Figure S2. Plant and fungal sterols support the development of *Ae. albopictus* to adulthood.** **A.** Complete statistical analysis of the data shown in Fig. 2A; *p*-values from Chi-square tests comparing each dietary sterol to EtOH control. **B.** Calibration curves for the different sterols used in the experiments. **C.** Representative GC–MS chromatograms of stigmasterol-fed larvae corresponding to Fig. 2I. **D.** Representative GC–MS chromatograms of larvae grown on ergosterol as the sole sterol source. **E.** GC–MS analysis of bacterial cultures without mosquitoes supplemented with cholesterol, stigmasterol, ergosterol, or ethanol (EtOH) control. The cultuers were analyzed after 14 days of incubation (n = 5 independent cultures per condition).

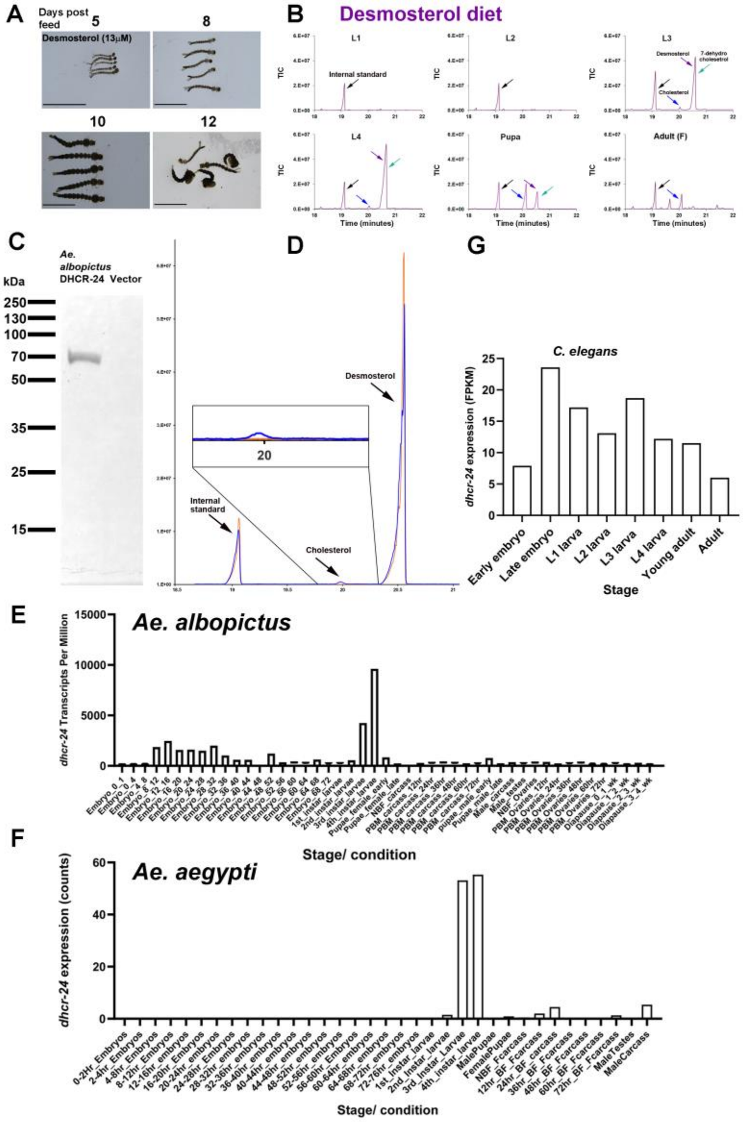


**Figure S3.** ***Ae. albopictus*** **DHCR-24 activity and regulation. A.** Representative images of larvae under desmosterol supplementation; anterior to the left; scale bars, 5 mm. **B.** Representative GC–MS chromatograms from desmosterol-fed larvae, corresponding to Fig. 3D. **C.** Western blot of bacterially expressed *Ae. albopictus* DHCR-24 tagged with a 6×His tag and detected using an anti-His antibody. **D.** GC–MS chromatograms of sterols extracted from lysates of bacteria expressing *Ae. albopictus* DHCR-24 and empty vector control. **E.** Comprehensive temporal RNA-seq analysis of *dhcr-24* expression in *Ae. albopictus*, using published developmental transcriptomic data[50]. NBM, no blood meal; PBM, post-blood meal. **F.** Comprehensive temporal RNA-seq analysis of *dhcr-24* expression in *Ae. aegypti*, using published developmental transcriptomic data[51]. NBM, no blood meal; BM, blood meal. **G.** Temporal RNA-sequencing analysis of *dhcr-24* expression in *C. elegans*, based on developmental expression data retrieved from WormBase.

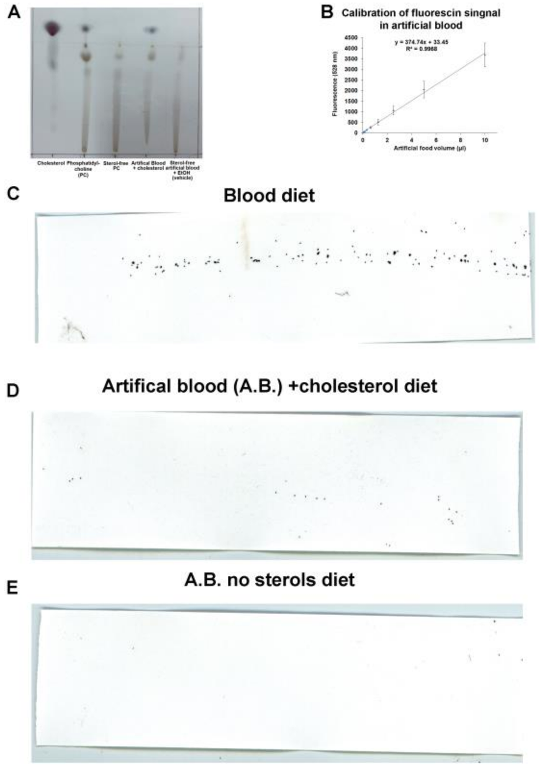


F

| Eggs per cage | 23.3.25 | 25.3.25 | 27.3.25 | 31.3.25 | Total/ repeat |
|---|---|---|---|---|---|
| Blood #1 | 43 | 165 | 20 | 140 | 368 |
| Blood #2 | 220 | 438 | 138 | 98 | 894 |
| Blood #3 | 105 | 395 | 107 | 34 | 641 |
| A.B.+ cholesterol #1 | 5 | 70 | 1 | 16 | 92 |
| A.B.+ cholesterol #2 | 28 | 0 | 1 | 0 | 29 |
| A.B.+ cholesterol #3 | 31 | 6 | 1 | 0 | 38 |
| A.B. no sterols #1 | 1 | 0 | 0 | 0 | 1 |
| A.B. no sterols #2 | 10 | 0 | 1 | 1 | 12 |
| A.B. no sterols #3 | 0 | 1 | 3 | 0 | 4 |

**Figure S4 | Cholesterol is essential for egg laying. A.** Thin-layer chromatography (TLC) analysis of artificial blood components before and after sterol extraction, demonstrating efficient sterol removal. **B.** Calibration curve of fluorescein in artificial blood; this curve was used to calculate the intake of artificial blood labeled with fluorescein in females (Fig. 4D). **C–E.** Representative images of egg laying in different cages under the indicated dietary conditions. **F.** Quantification of the number of eggs laid per cage per day in each independent replicate.
